# Comparative functional and evolutionary analysis of essential germline stem cell genes across the genus Drosophila and two outgroup species

**DOI:** 10.1101/2025.04.30.651540

**Authors:** Luke R. Arnce, Jaclyn E. Bubnell, Charles F. Aquadro

**Author notes:** First author and Correspondence: Luke R Arnce, Department of Molecular Biology and Genetics 233 Biotechnology Building, 526 Campus Rd. Ithaca, NY 14853.

## Abstract

In *Drosophila melanogaster, bag of marbles* (*bam*) encodes a protein essential for germline stem cell daughter (GSC) differentiation in early gametogenesis. Despite its essential role in *D. melanogaster*, direct functional evaluation of *bam* in other closely related *Drosophila* species reveal this essential function is not necessarily conserved. In *D. teissieri*, for example, *bam* is not essential for GSC daughter differentiation. Here, we generated *bam* null alleles using CRISPR-Cas9 in a species more distantly related to *D. melanogaster, D. americana*, to interrogate whether *bam*’s essential GSC differentiation function is novel to the *melanogaster* species group or a function more basal to the *Drosophila* genus. To further characterize the extent of the functional flexibility of other GSC regulating genes, we generated a gene ortholog dataset for 366 GSC regulating genes essential in *D. melanogaster* across 15 additional *Drosophila* and two outgroup species. We find that *bam*’s essential GSC function is conserved between *D. melanogaster* and *D. americana* and therefore originated prior to the formation of the *melanogaster* species group. Additionally, we find that ∼8% of the 366 GSC genes essential in *D. melanogaster* are absent in at least one of the 17 species in our ortholog dataset. These results indicate that developmental systems drift (DSD), in which the specific genes regulating a function may change, but the final phenotype is retained, occurs in stem cell regulation and the production of gametes across *Drosophila* species.

**Article summary:** Results from CRISPR induced *bam* null mutants in *D. americana* and comparative ortholog analysis of essential GSC regulating genes indicate that the evolutionary origin of *bam*’s essential GSC differentiation function is likely basal to the *Drosophila* genus, and there is functional flexibility in at least ∼8% of the 366 GSC regulating genes across the 17 included species.

## Introduction

Proper production of gametes is critical for reproduction, and in *Drosophila* it begins with the asymmetric division of germline stem cells (GSCs) to both self-renew, maintaining the germline, and the ultimate differentiation into sperm and eggs (Kahney et al. 2019).

Mis-regulation of this highly sensitive process can quickly lead to sterility, so coordination of these early cellular divisions might be presumed to be highly conserved (Gleason et al. 2018). However, recent results have demonstrated that several GSC regulating genes essential for reproduction in *D. melanogaster* show signs of positive selection through rapid amino acid diversification, are nonessential for fertility, and/or are completely absent in non-*melanogaster Drosophila* species (Civetta et al. 2006; Bauer DuMont et al. 2007; Bubnell et al. 2022), DuMont et al. 2021, Choi et al. 2015, Flores et al. 2015.

One of these GSC regulating genes is bag-of-marbles (*bam*). The encoded protein Bam is 442 amino acids with several known functions that are executed in complexes with other protein partners in *D. melanogaster.* The most well characterized of Bam’s functions is as the switch for GSC daughter differentiation, but Bam also has documented roles in the maintenance of gut integrity, and as a switch for preventing premature differentiation of hematopoietic progenitor cells (McKearin and Spradling 1990; Insco et al. 2009; Tokusumi et al. 2011).

*Bam* specifically acts as the switch gene for GSC differentiation for *D. melanogaster* females and is necessary for terminal differentiation of spermatogonia in males (McKearin and Spradling 1990). In females, *bam* is repressed in GSCs and bam expression causes differentiation by binding to several protein partners including *benign gonial cell neoplasm* (Bgcn) in order to repress the production of self-renewal factors Nanos and elF4a. The resultant differentiating cystoblast undergoes several mitotic divisions (Shen et al. 2009, Ohlstein et al. 2000, Li et al. 2013, Li et al. 2009).

Simultaneously, Bam concentrates at the fusome, which connects the cysts, and Bam and Bgcn function together to regulate the timing of mitotic divisions between cells. In males, Bam is expressed in GSCs, and as differentiation continues, expression increases (McKearin and Spradling 1990, Sgromo et al. 2018, Pan et al. 2014, Ji et al. 2017). Once *bam* expression reaches a threshold in the early spermatogonia, Bam binds to Bgcn and tumorous testis (*tut*) (Ting et al. 2013, Insco et al. 2009, Insco et al. 2012). Binding represses *mei-P26* and ends proliferation, triggering terminal differentiation and beginning meiosis (Shen et al. 2009, Ohlstein et al. 2000, Li et al. 2013, Li et al. 2009). Loss of *bam* function prevents differentiation in both sexes and causes over-proliferation of GSCs in females and spermatogonia in males, leading to tumors and sterility in *D. melanogaster* (Shivdasani et al. 2003, Ohlstein 1997).

Though *bam* plays an essential role in GSC regulation in *D. melanogaster*, recent results show *bam*’s sequence and function vary considerably across *Drosophila* or outgroup species (Bubnell et al. 2022). The total 442 amino acid Bam protein in *D. melanogaster* differs by 60 fixed amino acid differences (∼14%) with its sibling species *D. simulans*. Bam sequences differing from *D. melanogaster* by up to 308 (67%) of amino acids in other *Drosophila* species and up to 87% in outgroup species (Arnce, Bubnell, Aquadro 2025). Statistical tests of selection for *D. melanogaster* and *D. simulans bam* suggest that 94% and 72% of fixed amino acid differences respectively were driven by natural selection between the two current species and their common ancestor (Bubnell et al. 2022). Additional signals of positive selection at *bam* were detected in other *Drosophila* lineages across the genus leading to *D. yakuba*, *D. ananassae*, and *D. rubida* for example while other lineages, despite evaluation, showed no evidence of positive selection for protein diversification (Bubnell et al. 2022). This data suggests a striking level of sequence divergence, much of which potentially driven by natural selection, in orthologs of a gene critical for ensuring fertility in a *D. melanogaster*.

Additionally, recent functional studies of *bam* using complete loss-of-function (null) alleles in *Drosophila* species (Bubnell et al. 2022) revealed divergent roles across *D. simulans*, *D. melanogaster*, *D. teissieri*, *D. yakuba*, and *D. ananassae*. In *D. teissieri bam* null mutants showed no germline stem cell (GSC) differentiation defects in either sex, suggesting *bam* lacks its canonical role as a critical differentiation regulator in this species. While in *D. ananassae bam* null females were sterile, males exhibited normal spermatogenesis. These results demonstrate evolutionary divergence in *bam* sequence and function, despite its conserved essential role in GSC regulation in other *Drosophila* lineages.

Analyses of DNA polymorphism and divergence among other GSC regulatory genes within the *Drosophila melanogaster* species group have revealed distinct signatures of adaptive evolution across lineages (Bubnell et al. 2022). Notably, some core components of GSC regulatory networks exhibit dynamic evolutionary trajectories, including lineage-specific gene loss. For example, *Yb*—a gene expressed in somatic cap cells of the germarium and essential for GSC maintenance and transposable element suppression in *D. melanogaster* (Szakmary et al. 2009)—shows pronounced amino acid divergence between *D. melanogaster* and *D. simulans* (Flores et al. 2015). While *Yb* null mutants in *D. melanogaster* result in female sterility (Swan et al. 2001), preliminary evidence from our lab suggested that orthologs of *Yb* are absent in the *D. pseudoobscura, D. persimilis*, and *D. miranda* lineages raising the possibility of functional redundancy or network rewiring. These findings underscore both the evolutionary plasticity of GSC regulation and the potential for critical gene turnover within conserved developmental pathways.

A persistent challenge in comparative functional genetics lies in the widespread assumption of ortholog functional conservation, despite limited empirical validation. Most studies focus on single-species models, leaving ortholog activity across taxa largely inferred rather than experimentally confirmed (Tekaia 2016). Systematic assessment of the mechanistic basis and degree of functional conservation is crucial for reconstructing the evolution of gene networks. This limited validation of ortholog conservation introduces significant limitations when interpreting comparative genomic data: non-conserved ortholog functions imply divergent evolutionary pressures across lineages, thereby confounding hypotheses about ancestral genetic architectures or drivers of molecular evolution. Recent studies in bacteria and *Diptera* have begun to evaluate conservation of functional genes across closely related species and have identified surprising variability in ortholog functional conservation (Carranza et al. 2018, Bergmiller et al. 2012, Leeuwen et al. 2020, Deng et al. 2024, Hopkins et al. 2024, Zakerzade et al. 2025). Lineage-specific ortholog loss of functional reproductive genes has also been observed between humans and nonhuman primates (Carlisle et al. 2024). Altogether, these results suggest a more comprehensive comparative functional analysis of *bam* and other essential GSC regulating genes across the genus *Drosophila* could provide insight into the functional evolutionary history and extent of network flexibility of bam and essential GSC genes more broadly.

We executed our comparative functional analysis of *D. melanogaster*-essential GSC regulating genes using a two-pronged approach: generating a bam null allele in an additional divergent *Drosophila* species and evaluating ortholog presence or absence across 15 diverse *Drosophila* and two outgroup species (sheep blowfly *Lucillia cuprina* and house fly *Musca domestica*.

We chose to generate a *bam* null mutant *in D. americana* as this species represents a major, more divergent outgroup lineage to the *D. melanogaster* species group within the *Drosophila* genus and has previously been successfully edited with CRISPR/Cas9 (Vankuren et al. 2018, Lamb et al. 2020). We evaluated cytology and fertility in this null mutant using the strategy from Bubnell et al (2022) to evaluate *bam*’s function in GSC differentiation. This analysis adds broader evolutionary scope to our knowledge of *bam* function and provides additional insight into whether *bam*’s essential role in GSC differentiation is likely basal to all *Drosophila* species and was lost in specific lineages or whether *bam*’s critical role was a gained function within the *D. melanogaster* species group. Defining *bam* null phenotypes also provide a broader context for understanding the relationship between *bam* function and positive selection.

Null mutants are effective for performing comparative analyses of function for genes, like *bam*, with orthologs across species of interest. However, generating null mutants is expensive and time consuming in non-model species. Identifying ortholog absences in other species for GSC regulating genes essential in *D. melanogaster* provides an alternative rapid and cost-effective strategy for identifying potential functional differences in the genes and interaction networks that regulate germline stem cell maintenance, self-renewal and differentiation. To investigate ortholog functional conservation via ortholog presence or absence, we started with a set of 366 GSC regulating genes determined to be essential in *D. melanogaster* from a functional RNAi screen (Yan et al. 2014) along with two of *bam*’s close interacting partners that are also essential for fertility, *bgcn* and *Yb* (Szakmary et al. 2009, Li et al. 2009). We then used orthology tools in conjunction with a custom reciprocal blast best hit (RBBH) pipeline to identify ortholog presences/absences across 17 diverse *Drosophila* and two outgroup species. We also used the custom pipeline to identify physical and genetic GSC interacting gene orthologs and validate ortholog absences using a combination of localized PCR and sequencing when possible. Finally, we integrated functional categories as identified in Yan et al. (2014) for the 366 included GSC regulating genes. This generated gene ortholog dataset enables evaluation of the extent and characteristics of essential GSC regulating gene network flexibility as well as informed predictions regarding their functional evolutionary histories.

Here, we report that both female and male *D. americana bam* null mutants are sterile with GSC regulation defects, indicating *bam’*s essential GSC regulating function is not novel to the *melanogaster* species group, is likely basal to the genus Drosophila, and provides additional evidence that species in which *bam* is not necessary for gametogenesis (e.g., *D. teissieri* and *D. ananassae*) represent lineage-specific functional losses. Our comparative ortholog analysis of GSC regulating genes reveals that there is additional functional flexibility beyond *bam* with ∼8% (30 of 366) of the genes absent in one or more of the 17 included species and ∼3% absent in one or more of the 15 included *Drosophila* species. Surprisingly, we find that *Yb* is from *D. pseudoobscura* and *D. obscura* species, although it is necessary for GSC function and development, and therefore fertility, in *D. melanogaster*. Ortholog conservation does not necessarily indicate conservation of function (e.g., *bam*), so this represents the minimum functional flexibility in essential GSC regulating genes. These results altogether are consistent with other recent studies showing genes essential in one species for critical functions, like fertility, do not necessarily have the same essential function even among closely related species (Leeuwen et al. 2020). *Bam* functional variation and the absences of several essential GSC regulating gene across *Drosophila* are potential examples of developmental systems drift (DSD) or divergence in genetic systems that underpin a conserved phenotype (Weiss and Fullerton 2000, True and Haag 2001).

## Materials and methods

### Fly stocks and rearing

We raised fly stocks on standard cornmeal-molasses food at room temperature, and we used yeast-glucose food for fertility assays. We acquired lines with sequenced genomes for 15 *Drosophila* and two outgroup species: *Drosophila simulans* (strain: w501), *Drosophila sechellia* (strain: sech25), *Drosophila teissieri* (strain: GT53w), *Drosophila yakuba* (strain: Tai18E2)*, Drosophila takahashii* (strain: IR98-3 E-12201), *Drosophila elegans* (strain: 14027-0461.03)*, Drosophila serrata* (strain: Fors4), *Drosophila ananassae* (strain: 14024-0371.14) , *Drosophila pseudoobscura* (strain: MV2-25), *Drosophila obscura* (strain: BZ-5 IFL), *Drosophila willistoni* (strain: 14030-0811.24), *Drosophila mojavensis* (strain: 15081-1352.22), *Drosophila virilis* (strain: 15010-1051.87), *Drosophila grimshawi* (strain: 15287-2541.00), *Drosophila rubida* (strain: PH 161), *Musca domestica* (strain: aabys), *Lucilia cuprina* (strain: Lc7/37) (S1 Table A). We also acquired *D. simulans, sechellia, yakuba, serrata, willistoni, mojavensis, virilis,*and *grimshawi* from the National *Drosophila* Species Stock Center (NDSSC) (http://blogs.cornell.edu/drosophila/), *D. takahashii* from Kyorin-fly *Drosophila* species stock center (https://shigen.nig.ac.jp/fly/kyorin/), *D. teissieri*, *D. pseudoobscura*, the house fly *Musca domestica*, and the sheep blowfly *Lucillia cuprina* as gifts from Daniel Matute, Andy Clark, Jeffrey Scott, and Max Scott, respectively. *D. elegans* and *D. ananassae* were gifts from Artyom Kopp. *D. obscura* and *D. rubida* were gifts from Dmitri Petrov. We acquired the *D. americana* white eye mutant line (Lamb et al. 2020) for null generation as a gift from Trisha Wittkopp (S1 Table A).

### Strategy for generating a *bam* null phenotype in *D. americana*

We generated the *bam* null disruption by targeting the first exon of *bam* and introducing an early stop codon in the coding sequence using CRISPR/Cas9 gene editing. Because null homozygotes are sterile and thus cannot be maintained, we developed two *bam* disruption lines (one marked by 3x3P-Dsred and the other by 3xP3-YFP) which can be maintained as heterozygous lines (S1 Tables B-F). In order to phenotype the null homozygote, we crossed *bam*^3xP3-Dsred^/*bam^wt^* and *bam*^3xP3-YFP^/*bam^wt^*flies to create a *bam* disruption null homozygotes (*bam*^3xP3-Dsred^/*bam^YFP^*) which we then identified via fluorescent eye screen.

### Bam null construct cloning

Yasir Ahmed gifted us *bam* nucleotide sequences for *D. americana* (now on NCBI as G96 accession: PRJNA475270), and we performed cloning design in Geneious (S1 Table D). We generated PCR products using the NEB Q5 High Fidelity 2x master mix, then gel extracted and purified using the NEB Monarch DNA gel extraction kit. For PCR, sequencing, and cloning, we used IDT primers. We also generated donor plasmids for both the 3xP3-YFP and 3xP3-Dsred *bam* disruption lines using the strategy outlined in Bubnell et al. (2022). We prepared and purified plasmids for embryo injections with the Qiagen plasmid plus midi-prep kit followed by phenol-chloroform extraction for further RNase removal and then sequenced plasmids with whole plasmid sequencing (Plasmidsaurus).

### CRISPR/Cas9 and gRNA selection

We used Geneious to select gRNAs with no predicted off-targets in the reference genomes for *D. americana* (S3). We generated these as synthetic gRNAs (sgRNAs) from Synthego and used up to two gRNAs per injection to improve the likelihood of successful CRISPR events (S1 Table F).

### Embryo injections

Genetivision performed CRISPR/Cas9 injections including the appropriate plasmid donor, sgRNAs, and Cas9 protein (Synthego) into the *D. americana* line. We screened *bam* disruption lines for eye color to identify CRISPR/Cas9 mutant flies in-house using a Nightsea fluorescence system with YFP (cyan) and DsRed (green) filters. We backcrossed positive flies to generate lines which were maintained as heterozygous stocks. We confirmed CRISPR insertions by linear sequencing (Plasmidsaurus).

### Fertility assays

We executed following the strategy from Flores et al. (2015) for all female fertility assays. We collected and aged virgin females to sexual maturity (3-4 days for *D. americana*) (Markow and O’Grady 2006). We collected all generated genotypes from each bottle to control for bottle effects. Wildtype virgin males for each species were also aged until sexual maturity and distributed from different bottles across female genotypes. We crossed single females with two males, allowed to mate for nine days, then flipped onto new vials for nine more days, and then finally cleared from the vials while the offspring develop. We counted progeny daily and cleared to get total adult progeny per female. We also conducted male fertility assays with wildtype females and males of all generated genotypes. We executed fertility assays for *D. americana* on yeast-glucose food and all fertility experiments were kept at room temperature (approximately 21 degrees C).

### Fertility assay statistics

We used estimation statistics to assess fertility assay mean difference (effect size) in number of adult progeny between the wildtype *bam* genotype and the *bam* null homozygote and heterozygote genotypes. We generated estimation statistics and shared control Cumming plots using www.estimationstats.com (Ho 2019) (github.com/lukearnce/bam_null_ortholog). We used estimation statistics to enable determination of the size of the impact of the *bam* genotype on fertility via a non-parametric methodology. We reported significance as an effect size outside the 95% confidence interval (github.com/lukearnce/bam_null_ortholog).

### Immunostaining

We used the following primary antibodies: anti-Hts-1B1 (mouse, AB_528070, Developmental Studies Hybridoma Bank, concentrate 1:40) and anti-vasa (rat, AB_760351, DSHB, concentrate 1:20. We also used the following secondary antibodies: Alexaflour goat anti-rat 488 and goat anti-mouse 568 (Invitrogen) at 1:500. We performed immunostaining as described in Bubnell et al. (2022). In short, we digested ovaries and testes in cold 1x PBS and pipetted up and down to improve antibody permeability, fixed tissues in 4% paraformaldehyde, washed in PBST (1X PBS, 0.2% Triton-X 100), blocked in PBTA (1X PBS, 0.2% Triton-X 100, 3% BSA) (Alfa Aesar), and next, incubated in the appropriate primary antibody in PBTA overnight. We washed (PBST), blocked (PBTA), and incubated tissues in the appropriate secondary antibody for two hours. Tissue was then washed again (PBST) and finally mounted in mounting media with DAPI (Prolong glass antifade with NucBlue, Invitrogen) for imaging.

### Microscopy

We imaged ovaries and testes on a Zeiss i880 confocal microscope with 405, 488, and 568 nm laser lines at 40X (Plan-Apochromat 1.4 NA, oil) (Cornell BRC Imaging Core Facility). We analyzed and edited images using Fiji (ImageJ).

### McDonald-Kreitman test of selective neutrality for the *D. americana* lineage

We tested *D. americana bam* for significant departures from neutrality via McDonald-Kreitman test (MKT) (McDonald and Kreitman 1991) using polymorphism data gifted by Yasir Ahmed and the *bam* sequence from *D. virilis* as an outgroup. We implemented the strategy for MKT analysis of *D. americana bam* from Bubnell et al. (2022). In brief, we aligned *bam* sequences using PRANK (version) with the -codon and -F parameters using the PRANK tree guide. We used the codeml package from PAML (version 4.9) (Yang 1997, 2007) to generate the predicted common ancestor sequences for calculating lineage-specific divergence for *bam* with the MKT. We used PRANK alignments and trees as inputs to codeml with control file parameters (noisy=9, verbose=2, runmode=0, seqtype=1, CodonFreq=2, clock=0, aaDist=0, model=0, NSsites=0, icode=0, getSE=0, RateAncestor=1, Small_diff=.5e-6, cleandata=0, method=1) (github.com/lukearnce/bam_null_ortholog). We conducted an MKT comparing nonsynonymous and synonymous changes (Egea et al. 2008) using the http://mkt.uab.cat/mkt/mkt.asp webtool. We excluded polymorphic sites at less than 12% frequency, classified as slightly deleterious alleles not yet removed by purifying selection (Charlesworth and Eyre-Walker 2008). We used the predicted common ancestral species sequence to calculate lineage-specific divergence. We recorded values for the contingency table, the P-value of the χ^2^, alpha, and proportion of fixations predicted to be due to positive selection (Eyre-Walker 2006) (github.com/lukearnce/bam_null_ortholog).

### Genomes used in analysis

We downloaded genomes from NCBI for 15 *Drosophila* and two outgroup species: *Drosophila simulans* (assembly: Prin_Dsim_3.1), *Drosophila sechellia* (assembly: ASM438219v2), *Drosophila teissieri* (assembly: Prin_Dtei_1.1), *Drosophila yakuba* (assembly: Prin_Dyak_Tai18E2_2.1)*, Drosophila takahashii* (assembly: ASM1815269), *Drosophila elegans* (assembly: ASM1815250)*, Drosophila serrata* (assembly: Dser1.1), *Drosophila ananassae* (assembly: ASM1763931v2) , *Drosophila pseudoobscura* (assembly: UCI_Dpse_MV25), *Drosophila obscura* (assembly: ASM1815110v1), *Drosophila willistoni* (assembly: UCI_dwil_1.1), *Drosophila mojavensis* (assembly: ASM1815372v1), *Drosophila virilis* (assembly: Dvir_AGI_RSII-ME), *Drosophila grimshawi* (assembly: ASM1815329v1), *Drosophila rubida* (assembly: ASM3504616v1), *Musca domestica* (assembly: Musca_domestica.polishedcontigs.V.1.1), *Lucilia cuprina* (assembly: ASM2204524v1) (S1 Table A).

### Ensembl ortholog analysis

We used the Ensembl Compara online ortholog tool (Kersey et al. 2010 and Accessed date: June 2022) to collect ortholog predictions for 366 GSC regulating genes for the 10 *Drosophila* species that were available in Ensemble Compara (*D. melanogaster, D. simulans*, *D. sechellia*, *D. yakuba*, *D. ananassae*, *D. pseudoobscura*, D. willistoni, *D.* mojavensis, *D. virilis*, *D. grimshawi*) *and* two outgroup species (*L. cuprina* and *M. domestica*) (S2 Tables A-B). This tool catalogs relevant information about each ortholog including sequence alignment, target percent ID (percentage of orthologous sequence matching the *Drosophila melanogaster* sequence), query percent ID (percentage of *Drosophila melanogaster* sequence matching the orthologous sequence), gene order conservation score (evaluating synteny), and high or low ortholog confidence (calculated using results from other categories) (Kersey et al. 2010). Predicted orthologs were included as confident predictions if sequence alignment is equal to or greater than 25% identity (github.com/lukearnce/bam_null_ortholog).

### Reciprocal Best Blast Hit (RBBH) ortholog pipeline

To execute our RBBH ortholog analysis, implemented a multi-stage pipeline:

1. Initial RBBH

We first downloaded highly contiguous, long-read full genome sequences from NCBI for all included species (15 Drosophila species plus the two outgroup species). *D. teissieri, D. takahashii, D. elegans, D. serrata, D. obscura, D. rubida, M. domestica , and L. cuprina* were added to the initial set of species from Ensembl Compara. Then, we used custom scripts to perform forward and reverse BLASTp searches to identify potential orthologs and filter ortholog hits for GSC regulating genes as well as GSC gene interaction network genes (S2, github.com/lukearnce/bam_null_ortholog).

2. RBBH + syntenic evaluation

For genes with predicted absences, we conducted forward and reverse BLASTn searches as well as reciprocal BLASTp searches for the gene predicted absent and the syntenic genes (three on each side) flanking the GSC gene in *D. melanogaster*. We executed this to search for genes that are actually present but may have been predicted absent due to location at the end of contigs (S2, S3, github.com/lukearnce/bam_null_ortholog).

3. Direct validation of predicted gene ortholog absences

We evaluated predicted GSC gene ortholog absences with retained syntenic blocks directly via sequencing. We developed primers (IDT) for PCR amplification the sequence between retained syntenic genes. We then gel extracted, purified, and sequenced PCR products were to directly verify gene absence (S2, S3, github.com/lukearnce/bam_null_ortholog).

### Interaction networks and functional information

We cataloged physical interactors and genetic interactors from Flybase datasets for GSC regulating genes with predicted absences and evaluated interaction network genes for orthologs across included species using the same RBBH pipeline (Choi JY and Aquadro CF 2015) as well as ortholog predictions from Li et al. (2021). Annotated molecular functions (Choi and Aquadro 2015), functional categories identified by complex-enrichment analysis of the 366 GSC genes, and defect type, defined by the observed phenotypic effect of RNAi knockdown, were also incorporated from Yan et. al (2014). Defect types include GSC loss (cell viability), GSC loss (agametic), differentiation defect, and oocyte-specific phenotypes/late oogenesis (S3).

## Results

### Bam shows a lineage-specific signature of positive selection in *D. americana*

We detected a significant departure from neutrality suggesting positive selection favoring accelerated amino acid substitutions via the McDonald-Kreitman (1981) test for bam in the *D. americana lineage*. Contingency table values (Neutral Polymorphism: 42, Neutral Divergence: 0, Non-neutral Polymorphism: 27, and Non-neutral Divergence: 3) show the ratio of nonsynonymous to synonymous variation between species is greater than the same ratio within species, consistent with expectations for positive selection (χ^2^ = 4.390 P-value = 0.036) and 100% of fixations estimated to be due to positive selection (alpha = 1.00) (github.com/lukearnce/bam_null_ortholog). This raised the possibility that there may have been a functional change at *D. americana bam*.

### *Bam* is necessary for female and male fertility and germ cell differentiation in *D. americana*

We used CRISPR/Cas9 gene editing to generate *bam* null alleles in *D. americana*, a representative of a lineage divergent from *D. melanogaster* in the genus *Drosophila*. Wildtype (+/+) and *bam* null heterozygotes, (*bam*^3xP3-Dsred^/*bam^+^*) and (*bam*^3xP3-YFP^/*bam^+^*), are all fertile in both males and females (Fig. 1). However, *bam* null homozygotes, (*bam*^3xP3-Dsred^/*bam^3xP3-YFP^*), were completely sterile in males and females (P < 0.0001, permutation test, Fig. 1a&d) (github.com/lukearnce/bam_null_ortholog). As in *D. melanogaster, D. simulans, D. yakuba,* and *D. ananassae* (Bubnell, et al. 2022), one copy of wildtype *bam* is sufficient to rescue the *bam* null sterility phenotype in *D. americana*.

**Figure 1.**
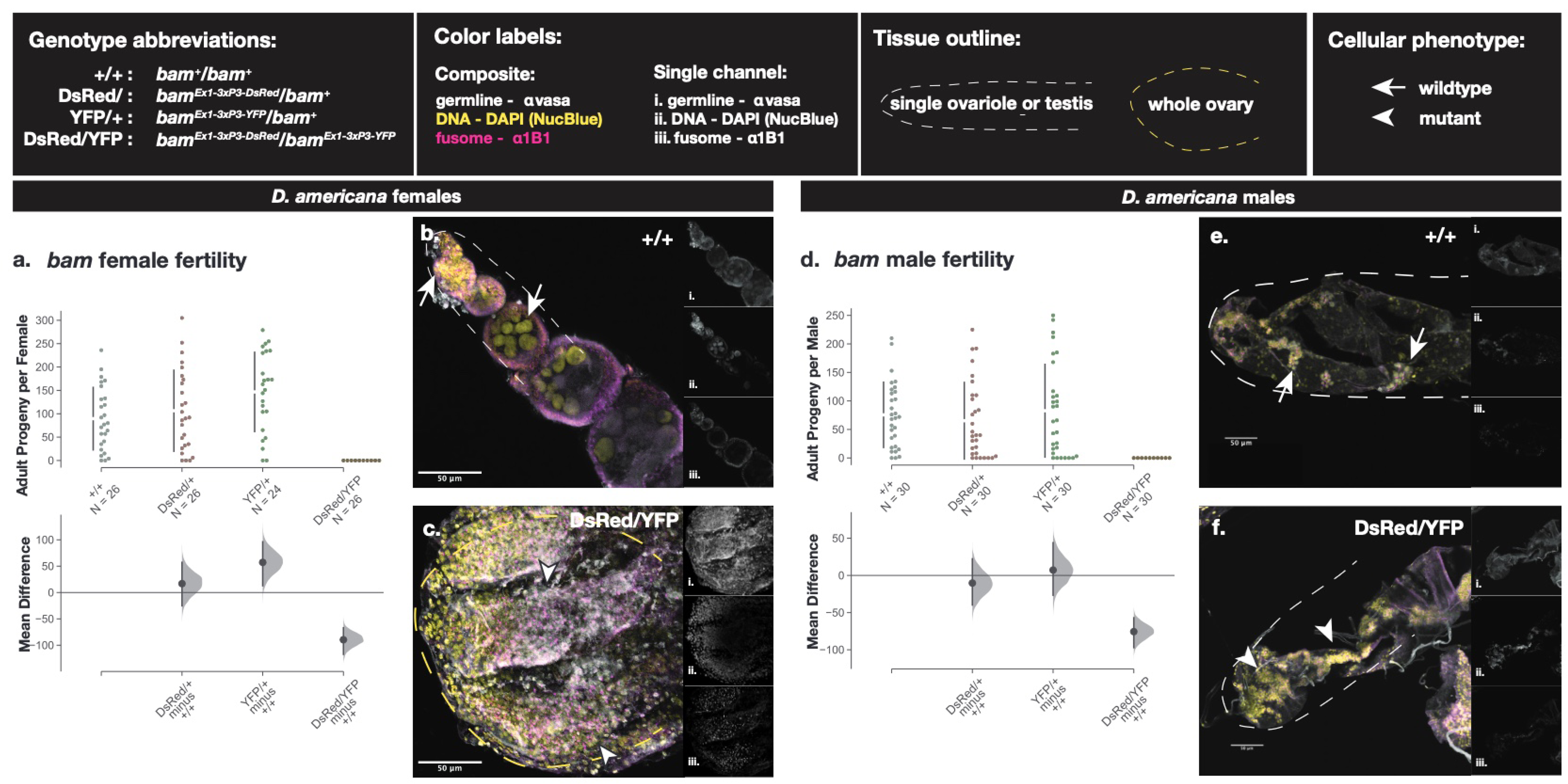
Fertility and cytological analyses of *bam* function in adult *D. americana.* Fertility is presented as adult *D. americana* progeny per fly for each *bam* genotype and presented separately for females (a) and males (d). The raw data as progeny per fly is plotted on the upper axes with the mean difference for the three genotype comparisons against the shared control wildtype illustrated in the Cumming estimation plots on the lower axes. Mean differences are plotted as bootstrap resampling distributions. Each mean difference is depicted as a dot, and 95% confidence intervals are indicated by the vertical black bars. Immunostaining of ovaries (b and c) and testes (e and f) of wildtype (b and e) and null *bam* genotypes (c and f). Composite Z-projections for ovaries and testes show staining for the germline (vasa), fusome (1B1), and nuclei (DAPI) with separate single channels for each image illustrated in the side panels (i. vasa, ii. DAPI, iii. 1B1). Wildtype tissue phenotypes are indicated with arrows and mutant tissue phenotypes are indicated with arrowheads.

**Figure 2.**
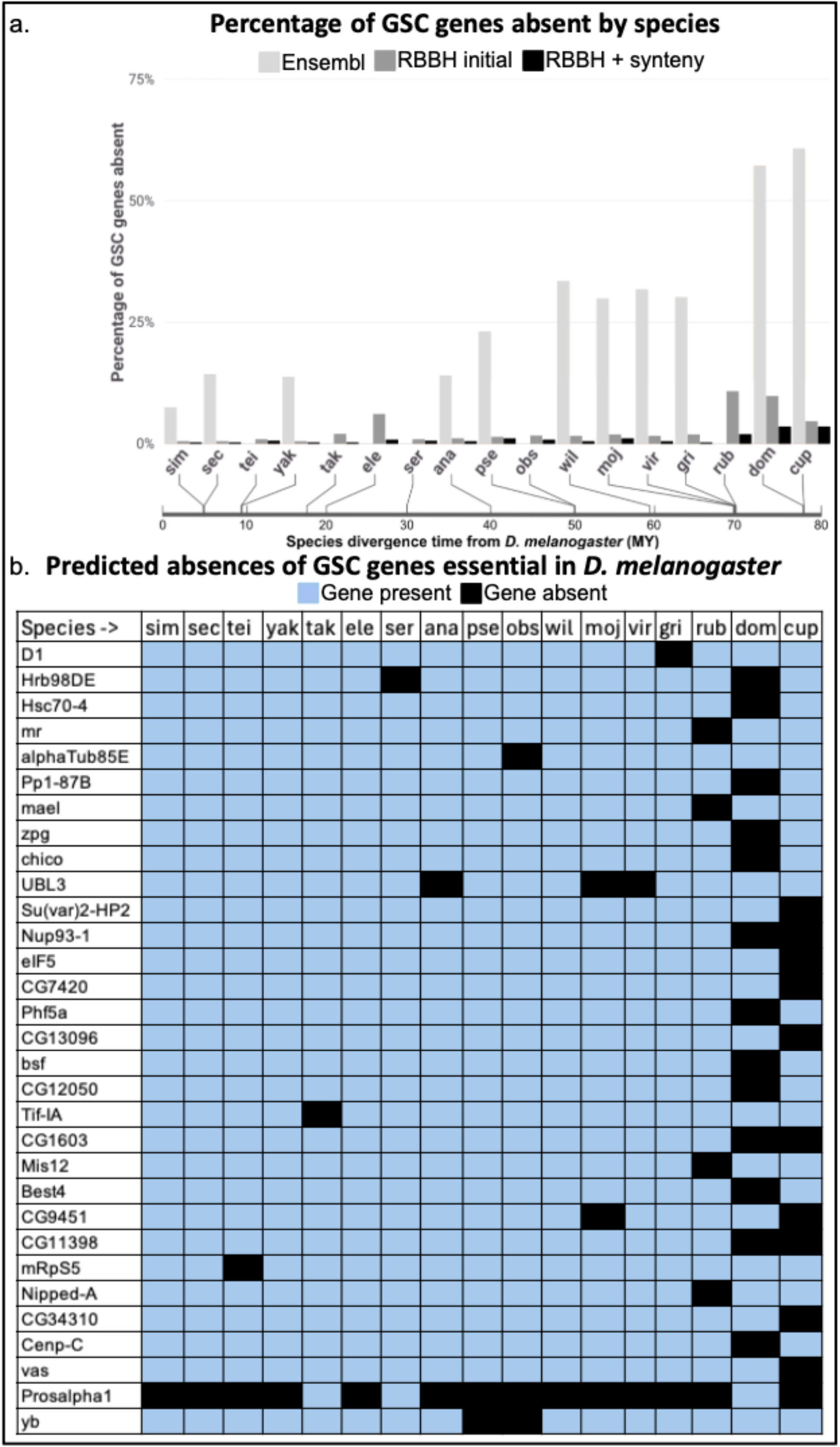
Predicted absences of 366 GSC regulating genes essential in *D. melanogaster* (predicted by Yan et al. 2014) by species plus *bgcn* and *Yb* using different ortholog detection strategies. a. Percentage of GSC genes predicted absent by species for three ortholog identification strategies. Light grey bars represent ortholog predictions using Ensembl, dark grey represent predictions with RBBH alone, and black represent RBBH plus syntenic evaluation. The bar below the species indicates divergence time from *D. melanogaster* (MY). b. Presence and absence of GSC genes across 15 *Drosophila* and two outgroup species that are predicted absent in at least one species after RBBH and syntenic evaluation. Blue indicates gene presence and black indicates gene absence.

To confirm the *bam* null sterile fertility phenotype was due to defects in GSC function as expected if *bam* function is conserved between *D. melanogaster* and *D. americana*, we evaluated the cytology of *D. americana bam* null ovaries and testes (Fig. 1). We imaged 3-5 day-old ovaries from *D. americana bam* wildtype and *bam* null females that we immunostained with antibodies to vasa and 1B1 and mounted with DAPI. (Lavoie et al. 1999). Homozygous *bam* null cytology recapitulated the classic *bag of marbles* phenotype, with over-proliferation of small, undifferentiated GSC-like cells in the ovaries (Fig. 1c) and testes (Fig. 1f) in contrast to *bam* wildtype ovaries (Fig 1b) and testes (Fig e) which consist of cysts made of larger differentiating germline cells. Our cytological data reveal that *bam* is necessary for early germ cell differentiation in *D. americana*, consistent with fertility assay results (Fig. 1a & d).

### Essential GSC gene ortholog absences across species

While functional genetic analyses for *bam* across diverse lineages in the genus *Drosophila* revealed some striking variation in *bam’s* role in fertility and germ cell differentiation, the financial and time costs required to generate null mutants for other GSC genes across diverse species are prohibitive at this point. Therefore, we next chose an alternative, albeit less sensitive, approach to evaluate the functional consistency of the roles of GSC regulating genes that are essential in *D. melanogaster*. Focusing on the experimentally defined set of GSC regulating genes determined to be essential in *D. melanogaster* (Yan et al. 2014), we defined a functional difference in GSC regulation pathways between species as the absence of an essential ortholog in *D. melanogaster* in any other *Drosophila* and/or outgroup species.

Our initial Ensembl ortholog analysis predicted absences for 311 of 366 GSC regulating genes for nine *Drosophila* and two outgroup species (S2 Tables A-B, github.com/lukearnce/bam_null_ortholog). Number of absences per species ranged from 27 (7.38%) in *D. simulans* to 222 (60.66%) in the outgroup species *Lucillia cuprina* (sheep blowfly). Next, we used our Reciprocal Best Blast Hit (RBBH) ortholog assessment that revealed predicted absences for 79 of 366 GSC regulating genes across 15 *Drosophila* and two outgroup species. Number of absences per species ranged from two (0.5%) in *D. simulans* to 39 (10.7%) in *D. rubida*. Finally, with our most stringent approach combining RBBH and syntenic evaluation of orthologs, we predicted absences for 30 of 366 GSC regulating genes across 15 *Drosophila* and two outgroup species (Fig. 3) (S3, github.com/lukearnce/bam_null_ortholog). Our most stringently assessed number of absences per species ranged from one (0.27%) in *D. simulans* to 13 (3.6%) in the outgroup species *Musca domestica* (house fly).

**Figure 3.**
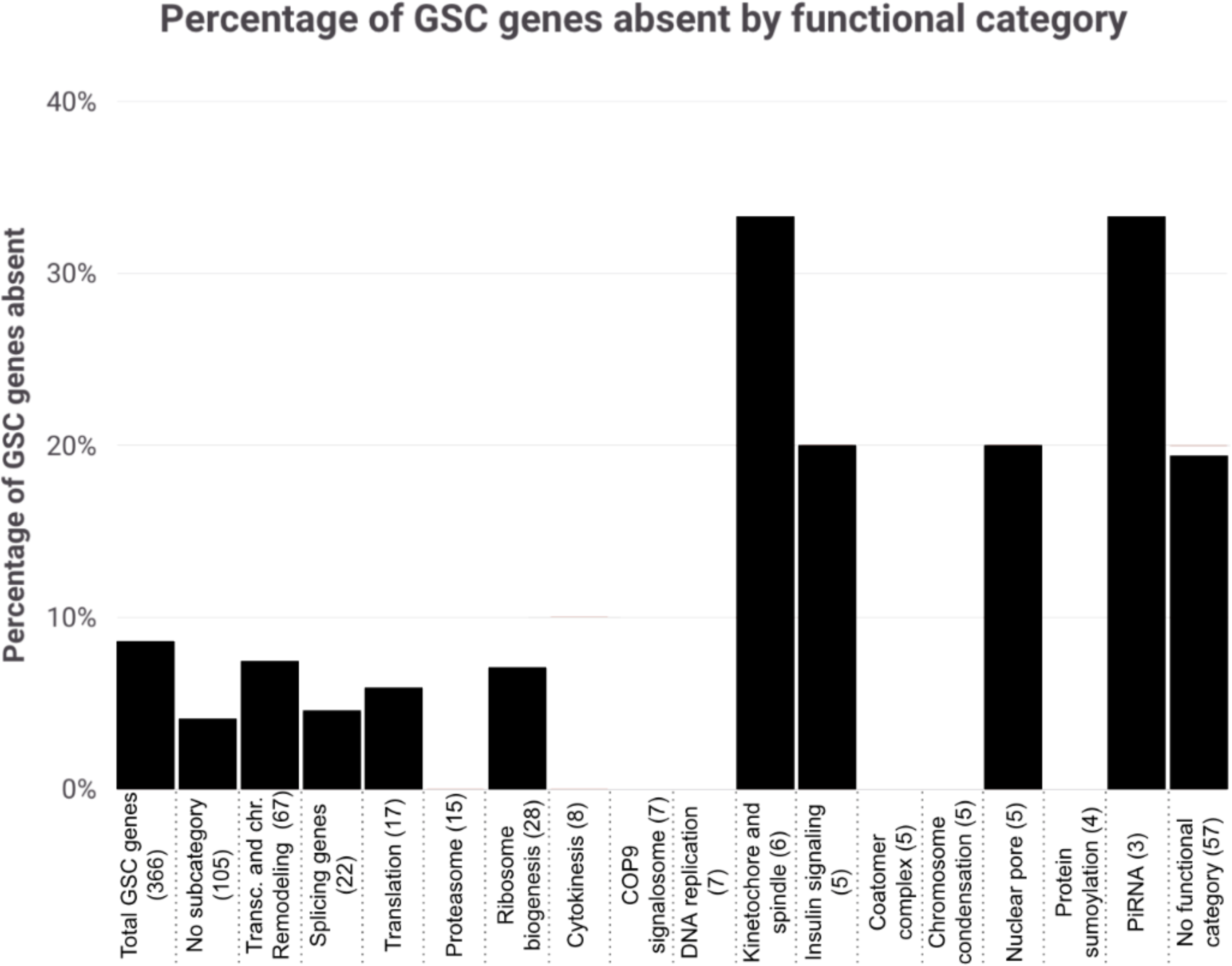
Percentage of GSC genes absent by functional category. Categories are pulled from the gene-interaction network generated in Yan et al. (2014). The “No subcategory (105)” group includes genes represented in the interaction network map without clear categorical associations and the “No functional category (57)” includes GSC genes that do not appear in the interaction network map.

### Verification of gene absences with retained syntenic blocks

To experimentally confirm the predicted GSC regulating gene ortholog absences we found with RBBH + syntenic evaluation with retained syntenic blocks, we PCR amplified and sequenced the syntenic block spanning the expected gene absence. The sequencing results verified all predicted absences with retained synteny therefore our RBBH + syntenic computational predictions represented real absences in the amplified regions (github.com/lukearnce/bam_null_ortholog). Using our final ortholog presence and absence results, we then sought to identify potential patterns in gene presence or absence across the included species. Primarily, we evaluated whether and in what ways essential gene conservation varied across gene functional categories and interaction network size (S3).

### Variable GSC gene conservation across species and functional categories

We found that GSC regulating genes with predicted absences are unequally represented across functional categories identified by complex enrichment. At the extremes, there are zero predicted absences in the proteasome functional category (15 genes) and two of six genes (33%) in the Kinetochore and spindle functional category (Fig. 3). Absences categorized by defect types show less variation ranging from 7.14% for GSC loss (cell viability) (12 of 168) to 11.11% for genes with oocyte-specific phenotypes/late oogenesis (five of 45).

### Variability in absent GSC gene interaction network size and absences

Of the 30 GSC regulating genes (from the 366 in Yan et al. (2014)) with predicted absences across included species, 23 genes have physical and/or genetic interactions (Fig. 4, S3). 13 of these 23 genes also have predicted absences in their interaction networks, and most of these interaction network absences are in the same species as their related absent GSC regulating gene (Fig. 5, S3).

**Figure 4.**
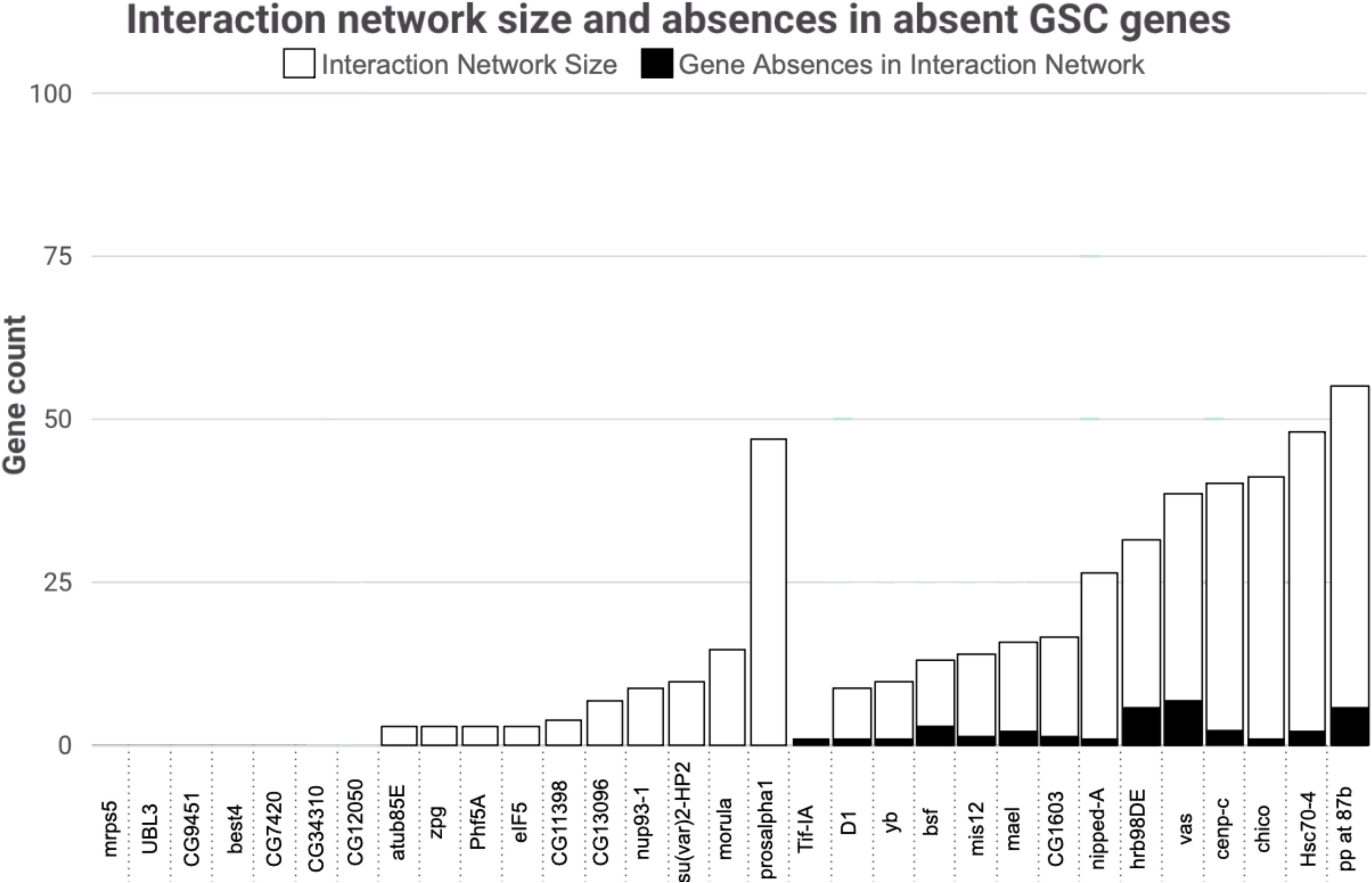
Interaction network size and absences in absent GSC genes. Genes with at least one absence in the included species are listed with white bars representing the size of their interaction networks including genetic and physical interactors. The number of interaction network genes absent are represented with black bars.

**Figure 5.**
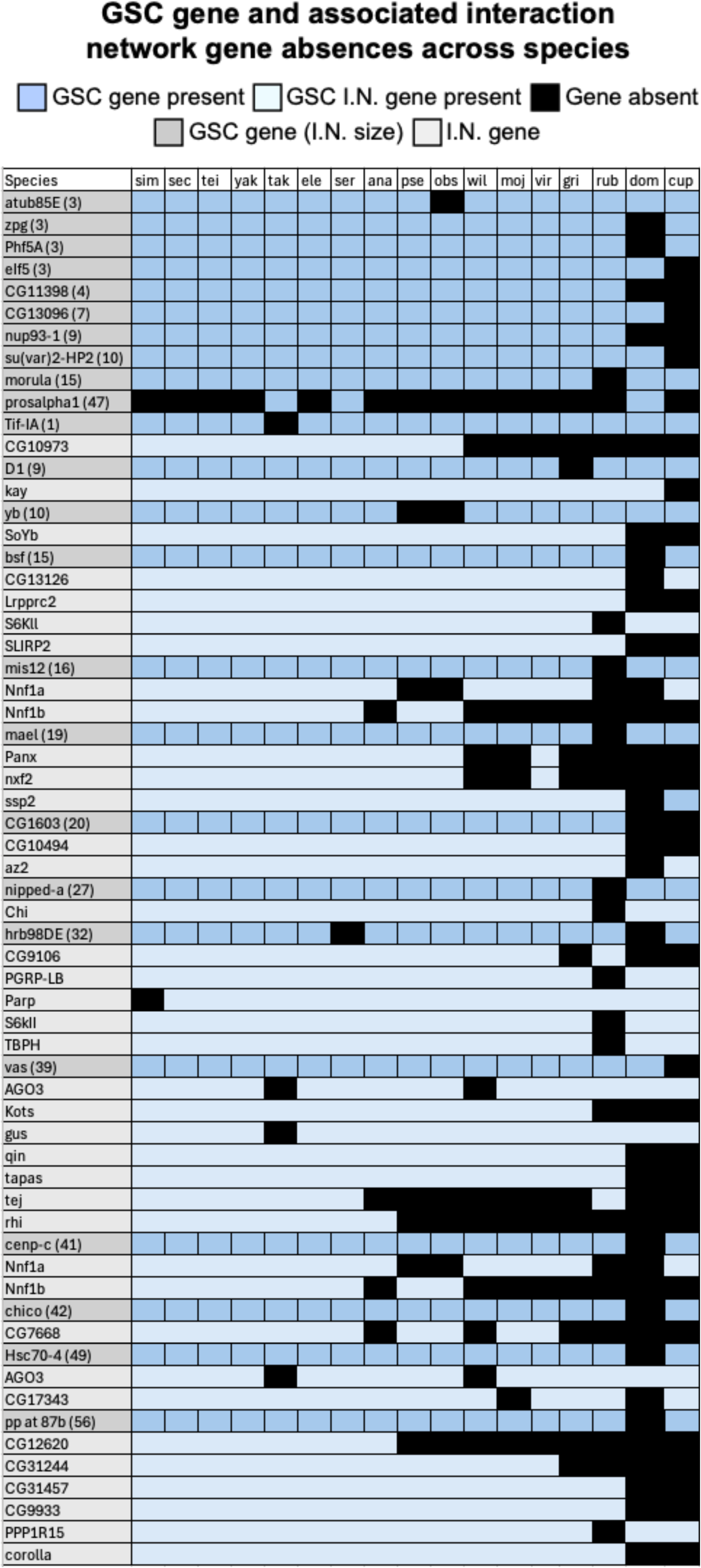
GSC gene and associated network absences across species. GSC genes with absences in at least one included species are highlighted in dark grey. Associated interaction network (I.N.) genes are indented, italicized, and highlighted in light grey. Gene presence is indicated by light blue and absence is indicated by black. First, GSC genes with no absences in their interaction networks are arranged by increasing interaction network size. Next, GSC genes with absences in their interaction networks are arranged in the same manner. GSC genes with no interaction networks (six genes) are excluded from this figure.

## Discussion

Fertility and germline cytological assays of *bam* null mutants in *D. americana* reveal that *bam* is essential for GSC differentiation in both males and females. This demonstrates that *bam*’s essential function in GSC differentiation is not novel to the *D. melanogaster* species group, and its evolutionary origin in this function likely occurred just prior to or just after the origin of the genus Drosophila. *Bam* functional differences in *D. teissieri* and *D. ananassae* likely represent lineage-specific GSC functional losses. Similar losses may exist among the many other non-tested taxa in this species-rich genus.

These analyses of *bam* function show that high amino acid sequence variability between species does not necessarily imply functional divergence. *D. americana bam* and *D. melanogaster bam* share only 35% sequence identity, yet both execute the same essential function in GSC daughter differentiation. In contrast, *D. teissieri bam* and *D. melanogaster bam* share 75% sequence identity and they are functionally distinct.

Conservation of sequence does not necessarily imply conservation of function and divergence of sequence does not necessarily imply a divergence in function. A classic example of this dynamic includes cytochrome c (Garrido et al. 2006). Cytochrome c is one of the most conserved proteins among eukaryotes with an amino acid sequence conservation of 70-90% between species as divergent as yeast and mammals. In many yeasts and lower eukaryotes, cytochrome c functions solely as an electron carrier in the mitochondrial respiratory chain while in mammals the same (or nearly identical) cytochrome c also acquired an additional role as an apoptotic signal by helping to trigger caspase activation and programmed cell death. This extra “moonlighting” function in apoptosis is a dramatic divergence in biological role despite a highly conserved structure (Garrido et al. 2006).

The resolution and depth of our analysis was made possible due to the large collection and phylogenetic density of high-quality genomes for a growing number of species in the genus *Drosophila* in combination with the depth of functionally defined genes within *D. melanogaster*. Large-scale gene ortholog identification is mostly not possible in species with only low-quality genomes available, and, potentially as a result, there are currently very few large-scale comparative analyses of gene orthologs, essential or nonessential (Carranza et al. 2018, Bergmiller et al. 2012, Leeuwen et al. 2020, Deng et al. 2024). However, as more high-quality genomes are produced and published for more species, this type of comparative ortholog analysis becomes more feasible. Even when using only species with available high-quality genomes for comparative analysis, efforts must be made to ensure potential orthologs and absences are properly identified. For all of the GSC regulating genes in *D. melanogaster* that we analyzed (366 from Yan et al. (2014) plus *bgcn* and *Yb*), ortholog predictions vary dramatically based on identification strategy. Using Ensembl alone, we predicted 85% (311 of 366) of GSC regulating genes absent in at least one included species while we only predicted ∼8% (30 of 366) absent based on the more detailed evaluation incorporating RBBH and synteny. Due to the significant discrepancy in predictions, the predicted presence or absence of orthologs via Ensembl must be further interrogated by additional strategies including RBBH and syntenic evaluation before confidently characterizing predictions as orthologs that are truly absent. As an example, benign gonial cell neoplasm (*bgcn*) is crucial gene that is broadly conserved, including a human ortholog, that was predicted to be absent in the initial RBBH evaluation. However, using follow-up BLASTn and syntenic evaluation we found the *bgcn* ortholog split over short contigs that made initial identification difficult. In contrast, *Yb*, an additional essential GSC regulating gene in *D. melanogaster* was computationally predicted absent in *D. obscura* and *D. pseudoobscura*. Follow-up syntenic evaluation and sequencing confirmed these absences of *Yb* from their syntenic regions in these two species.

The 30 GSC (out of 366) GSC regulating genes that are confidently predicted to be absent represent significant flexibility in essential GSC regulating genes beyond *bam* across species. These results add to a growing body of evidence that suggests the general ortholog assumption of shared function is not universal and provide examples of developmental systems drift (DSD) in a process critical for early reproduction. These 30 genes are also not equally distributed across functional categories. While there are no predicted absences in proteasome genes, two of six (33%) genes involved in the kinetochore and spindle have predicted absences. This variable flexibility across functional categories suggests there may be some networks that are more flexible in their regulation while others are particularly intolerant. Additionally, further analysis of these 30 genes with absent orthologs shows that these genes generally have interaction networks including genetic and physical interactors. Absences in orthologs of interaction network genes also generally occur in the same species that their related GSC gene is absent. This could potentially indicate that having a network of interacting partners more easily enables gene loss by compensating for the function of the absent gene through redundancy or alternative regulation while essential genes without large interaction networks could generally be more difficult to replace (Albalat 2016, Wagner 2008, Carroll 2008). One gene, *Prosalpha1*, is absent in the most species of the included GSC genes. In addition to having a large interaction network (47 genes), it also has a very similar gene, *Prosalpha1R*, within the surrounding syntenic region. *Prosalpha1R* could possibly facilitate alternative execution of *Prosalpha1’*s essential function.

Our findings here provide additional evidence that orthologous genes do not necessarily function identically even when critical for regulating essential systems. Despite the essential nature of proper gamete development, what has been described as developmental systems drift (DSD) is occurring in genes critical for regulating the earliest stages of this process across *Drosophila* and closely related outgroups.

Spermatogenesis regulating genes further downstream of GSC regulation have also been evaluated across some *Drosophila* species. Results showcase similar functional flexibility with duplications or losses of sperm nuclear basic proteins as well as the emergence of many functional *de novo* genes (Lee U 2025, Chang C 2023). Some of the particular genes involved in the process change, but the final phenotype is maintained across species.

We conclude that ortholog function must be evaluated in a species-specific manner. These 30 GSC genes and *Yb* that show absences in some species represent the baseline of functional flexibility for the *D. melanogaster* essential GSC regulating genes, but many other GSC regulating genes still present, like *bam*, could have variable function across species. The extent of developmental systems drift occurring beyond GSC regulating genes in these 17 species is largely unknown, but further evaluation of ortholog functional flexibility focusing on different organisms and phenotypes could reveal differences in the key genes regulating any number of traits across species.

Results from this dataset suggest some potential patterns related to ortholog loss of essential genes, but more extensive analysis needs to be executed to make more concrete assessments of the prevalence and characteristics of ortholog functional flexibility.

## Data Availability

All raw and filtered data are available in a public repository at www.github.com/lukearnce/bam_null_ortholog. These data include the following:

### McDonald-Kreitman test for *D. americana bam*

PRANK inputs and control parameters for the *D. americana bam* MK test with contingency table values, the P-value of the χ^2^, alpha, and proportion of fixations predicted to be due to positive selection.

### *D. americana bam* CRISPR Null fertility assays

D. americana male and female estimation statistics and significance as an effect size outside of the 95% confidence intervals

### GSC ortholog identification

Initial RBBH ortholog analysis results, Full genomes, custom scripts to execute forward and reverse BLASTp, filtered ortholog hits, interaction network genes, RBBH with syntenic evaluation gene hits, and sequence verification of predicted ortholog absences

## Acknowledgements

We thank Jolie Carlisle for her thoughtful comments on an earlier version of the manuscript. This work was supported by National Institute of Health (United States) R01-GM095793 to Charles F. Aquadro.

## References

Albalat R, Canestro C 2016 Evolution by gene loss. Nature Reviews Genetics 17, 379–391. 10.1038/nrg.2016.39

Arnce LR, Bubnell JE, Aquadro CF 2025 Comparative Analysis of *Drosophila* Bam and Bgcn Sequences and Predicted Protein Structural evolution. Journal of Molecular Evolution 10.1101/2024.12.17.628990

Bauer DuMont et al. 2007 Recurrent Positive Selection at Bgcn, a Key Determinant of Germ Line Differentiation, Does Not Appear to be Driven by Simple Coevolution with Its Partner Protein Bam. Molecular Biology and Evolution, 24(1):1882–191. 10.1093/molbev/msl141

Bergmiller T et al. 2012 Patterns of Evolutionary Conservation of Essential Genes Correlate with Their Compensability. PLOS Genetics, 10.1371/journal.pgen.1002803

Carlisle J et al. 2024 Recurrent Independent Pseudogenization Events of the Sperm Fertilization Gene ZP3r in Apes and Monkeys. Journal of Molecular Evolution, 92:695–702. 10.1007/s00239-024-10192-x

Carranza S et al. 2018 Diversity, distribution and conservation of the terrestrial reptiles of Oman (Sauropsida, Squamata). PLOS One, 10.1371/journal.pone.0190389

Carroll, S. 2008 Evo-devo and an expanding evolutionary synthesis: a genetic theory of morphological evolution. Cell 134(1)25–36. 10.1016/j.cell.2008.06.030

Chang C et al. 2023 Expansion and loss of sperm nuclear basic protein genes in *Drosophila* correspond with genetic conflicts between sex chromosomes. eLife, 10.75554/eLife.85249

Charlesworth J and Eyre-walker A 2008 The McDonald-Kreitman test and slightly deleterious mutations. Molecular Biology and Evolution, 25(6):1007–15. 10.1093/molbev/msn005

Choi JY and Aquadro CF 2015 Molecular Evolution of Drosophila Germline Stem Cell and Neural Stem Cell Regulating Genes. Genome Biology and Evolution, 7(11):3097–114. 10.1093/gbe/evv207

Civetta A et al. 2006 Rapid evolution and gene-specific patterns of selection for three genes of spermatogenesis in Drosophila. Molecular Biology and Evolution, 23(3):655–62. 10.1093/molbev/msj074

Deng D et al. 2024 Functional Divergence in Orthologous Transcription Factors: Insights from AtCBF2/3/1/ and OsDREB1C Molecular Biology and Evolution, 41(5):msae089. 10.1093/molbev/msae089

DuMont, V.L., White, S.L., Zinshteyn, D. and Aquadro, C.F. 2021. Molecular population genetics of *Sex-lethal (Sxl)* in the *D. melanogaster* species group - a locus that genetically interacts with *Wolbachia pipientis* in *Drosophila melanogaster*. G3 Genes|Genomes|Genetics 11(8), jkab197. 10.1093/g3journal/jkab197

Egea R et al. 2008 Standard and generalized McDonald-Kreitman test: a website to detect selection by comparing different classes of DNA sites. Nucleic Acids Research, 36: 157–162. 10.1093/nar/gkn337

Eyre-Walker A et al. 2006 The distribution of fitness effects of new deleterious amino acid mutations in humans. Genetics, 173(2):891–900. 10.1534/genetics.106.057570

Flores HA, Bubnell JE, Aquadro CF, Barbash DA. 2015 The Drosophila bag of marbles Gene Interacts Genetically with Wolbachia and Shows Female-Specific Effects of Divergence. PLOS Genetics 185(4):613–27. 10.1371/journal.pgen.1005453

Garrido C et al. 2006 Mechanisms of cytochrome c release from mitochondria. Cell Death & Differentiation 13, 1423–1433. 10.1038/sj.cdd.4401950

Gleason R et al. 2018 Protecting and Diversifying the Germline. Genetics, 208(2):435–471. 10.1534/genetics.117.300208

Ho J et al. 2019 Moving beyond P values: data analysis with estimation graphics. Nature Methods, 16(7):565–566. 10.1038/s41592-019-0470-3

Hopkins B, et al. Decoupled evolution of the *Sex Peptide* gene family and *Sex Peptide Receptor* in Drosophilidae. PNAS 121(3):e2312380120. 10.1073/pnas.2312380120

Insco ML, Leon A, Tam CH, McKearin DM, Fuller MT. 2009 Accumulation of a differentiation regulator specifies transit amplifying division number in an adult stem cell lineage. Proc Natl Acad Sci U S A. 106(52) 22311–22316. 10.1073/pnas.0912454106

Insco ML, Bailey AS, Kim J, Olivares GH, Wapinski OL, Tam CH, et al. 2012 A self-limiting switch based on translational control regulates the transition from proliferation to differentiation in an adult stem cell lineage. Cell Stem Cell. 11: 689–700. 10.1016/j.stem.2012.08.012

Ji S, Li C, Hu L, Liu K, Mei J, Luo Y, et al. 2017 Bam-dependent deubiquitinase complex can disrupt germ-line stem cell maintenance by targeting cyclin A. Proc Natl Acad Sci U S A. 114(24):6316–6321. 10.1073/pnas.1619188114

Kahney E et al. 2019 Regulation of *Drosophila* germline stem cells. Current Opinion in Cell Biology, Vol. 60, 27–35. 10.1016/j.ceb.2019.03.008

Kersey PJ et al. 2010 Ensembl Genomes: Extending Ensembl across the taxonomic space Nucleic Acids Research Vol. 38 D563-D569. 10.1093/nar/gkp871

Lamb AM, Wang Z, Simmer P, Chung H, Wittkopp P 2020 *ebony* Affects Pigmentation Divergence and Cuticular Hydrocarbons in *Drosophila americana* and *D. novamexicana* Front. Ecol. Evol. 10.3389/fevo.2020.00184

Lee U et al. 2025 Comparative single cell analysis of transcriptional bursting reveals the role of genome organization on *de novo* transcript origination. bioRxiv preprint.

Leeuwen J et al. 2020 Systematic analysis of bypass suppression of essential genes. Mol Syst Biol, 16:1–24. 10.15252/msb.20209828

Li F et al. 2021 Phylogenomic analyses of the genus *Drosophila* reveals genomic signals of climate adaptation. Molecular Ecology Resources. 10.1111/1755-0998.13561

Li Y, Minor NT, Park JK, McKearin DM, Maines JZ. 2009 Bam and Bgcn antagonize Nanos-dependent germ-line stem cell maintenance. Proc Natl Acad Sci U S A. 106: 9304–9309. 10.1073/pnas.0901452106

Li Y, Zhang Q, Carreira-Rosario A, Maines JZ, Mckearin DM. 2013 Mei-P26 Cooperates with Bam, Bgcn and Sxl to Promote Early Germline Development in the Drosophila Ovary. PLoS One. 8: 58301. 10.1371/journal.pone.0058301

Markow T and O’Grady P 2006 Drosophila: A Guide to Species Identification and Use. Book

McDonald J and Kreitman M 1991 Adaptive protein evolution at the Adh locus in Drosophila Nature 351(6328):652-4. 10.1038/351652a0

McKearin DM, Spradling AC. 1990 Bag-of-marbles: A Drosophila gene required to initiate both male and female gametogenesis. Genes Dev. 4: 2242–2251. 10.1101/gad.4.12b.2242

Ohlstein B, Lavoie CA, Vef O, Gateff E, Mckearin DM. 2000 The Drosophila Cystoblast Differentiation Factor, benign gonial cell neoplasm, Is Related to DExH-box Proteins and Interacts Genetically With bag-of-marbles. 155(4):1809–19. 10.1093/genetics/155.4.1809

Ohlstein B, McKearin D. 1997 Ectopic expression of the Drosophila Bam protein eliminates oogenic germline stem cells. Development. 124(18):3651–62. 10.1242/dev.124.18.3651

Pan L, Wang S, Lu T, Weng C, Song X, Park JK, et al. 2014 Protein competition switches the function of COP9 from self-renewal to differentiation. Nature. 514: 233–236. 10.1038/nature13562

Sgromo A, Raisch T, Backhaus C, Keskeny C, Alva V, Weichenrieder O, et al. 2018 Drosophila Bag-of-marbles directly interacts with the CAF40 subunit of the CCR4–NOT complex to elicit repression of mRNA targets. Rna. 24: 381–395. 10.1261/rna.064584.117

Shen R, Weng C, Yu J, Xie T. 2009 eIF4A controls germline stem cell self-renewal by directly inhibiting BAM function in the Drosophila ovary. Proc Natl Acad Sci U S A. 106: 11623–11628. 10.1073/pnas.0903325106

Shivdasani AA, Ingham PW. 2003 Regulation of Stem Cell Maintenance and Transit Amplifying Cell Proliferation by TGF-β Signaling in Drosophila Spermatogenesis. Curr Biol. 13(23):2065–72. 10.1016/j.cub.2003.10.063

Swan A, Hijal S, Hilfiker A, Suter B 2001 Identification of new X-chromosomal genes required for Drosophila oogenesis and novel roles for fs(1)Yb, brainiac and dunce Genome Res. 11(1):67–77. 10.1101/gr.156001

Szakmary A, Reedy M, Qi H, Lin H 2009 The Yb protein defines a novel organelle and regulates male germline stem cell self-renewal in *Drosophila melanogaster* J Cell Biol. 185(4): 613–627. 10.1083/jcb.200903034

Tekaia F 2016 Inferring Orthologs: Open Questions and Perspectives. Genomics Insights 9:17–28. 10.4137/GEI.S37925

Ting X 2013 Control of germline stem cell self-renewal and differentiation in the Drosophila ovary: concerted actions of niche signals and intrinsic factors. Wiley Interdiscip Rev Dev Biol. 2: 261–273. 10.1002/wdev.60

Tokusumi T. et al. 2011 Germ line differentiation factor Bag of Marbles is a regulator of hematopoietic progenitor maintenance during Drosophila hematopoiesis. Development, 138(18):3879–84. 10.1242/dev.069336

True JR, Haag ES. 2001. Developmental system drift and flexibility in evolutionary trajectories. Evol Dev. 3(2):109–119. 10.1046/j.1525-142x.2001.003002109.x

Vankuren NW, Long M. 2018 Gene duplicates resolving sexual conflict rapidly evolved essential gametogenesis functions. Nature Ecology and Evolution 2, 705–712. 10.1038/s41559-018-0471-0

Wagner A. 2008 Robustness and evolvability: a paradox resolved. Proc Biol Sci 275(1630):91–100. 10.1098/rspb.2007.1137

Weiss KM, Fullerton SM. 2000. Phenogenetic drift and the evolution of genotype-phenotype relationships. Theor Popul Biol. 57(3):187–195. 10.1006/tpbi.2000.1460

Yan D et al. 2014 A Regulatory Network of *Drosophila* Germline Stem Cell Self-Renewal Developmental Cell Vol 28 Issue 4 p. 459–473. 10.1016/j.devcel.2014.01.020

Zakerzade R, et al. 2025 Diversification and recurrent adaptation of the synaptonemal complex in *Drosophila*. PLoS Genetics 21(1):e1011549. 10.1371/journal.pgen.1011549

